# Kinesin-II motors differentially impact biogenesis of distinct extracellular vesicle subpopulations shed from *C. elegans* sensory cilia

**DOI:** 10.1101/2021.12.19.473369

**Authors:** Michael Clupper, Rachael Gill, Malek Elsayyid, Denis Touroutine, Jeffrey L. Caplan, Jessica E. Tanis

## Abstract

Extracellular vesicles (EVs) are bioactive lipid-bilayer enclosed particles released from nearly all cells. One specialized site for EV shedding is the primary cilium, a conserved signaling organelle. The mechanisms underlying cargo enrichment and biogenesis of heterogeneous EVs shed from cilia are unclear. Here we discover the conserved ion channel CLHM-1 as a new ciliary EV cargo. Using super-resolution microscopy, we imaged EVs released into the environment from sensory neuron cilia of *C. elegans* expressing fluorescently-tagged CLHM-1 and TRP polycystin-2 channel PKD-2 EV cargoes at endogenous levels. We find that these proteins are enriched in distinct EV subpopulations that are differentially shed in response to availability of hermaphrodite mating partners. Both CLHM-1 and PKD-2 localize to the ciliary base and middle segment of the cilium proper, but PKD-2 alone is present in the cilium distal tip and EVs shed from this site. CLHM-1 EVs released into the environment bud from a secondary site, the periciliary membrane compartment at the ciliary base. We show that individual heterotrimeric and homomeric kinesin-II motors have discrete impacts on the colocalization of PKD-2 and CLHM-1 in both cilia and EVs. Total loss of kinesin-II activity significantly decreases shedding of PKD-2 but not CLHM-1 EVs. Our data demonstrate that anterograde kinesin-II-dependent intraflagellar transport is required for selective enrichment of specific protein cargoes into heterogeneous EVs with different signaling potentials.

## INTRODUCTION

Nearly all cells release extracellular vesicles (EVs), which mediate the intercellular transport of biological macromolecules. These lipid bilayer-enclosed vesicles are typically categorized as either exosomes, which arise from the endosomal recycling pathway, or ectosomes, which are shed directly from the plasma membrane^1,2^. Dictated by cell of origin, EV cargoes play active roles in regulating physiological processes and propagating pathological conditions^3–5^. A single cell can release multiple discrete subpopulations of both exosomes and ectosomes, which are enriched with different molecular cargoes and have distinct physiological functions^6,7^. Questions remain regarding how cells enrich specific cargoes into EVs in order to foster this heterogeneity.

Bioactive EVs are shed from primary cilia^8–11^, highly conserved microtubule-based organelles that protrude from most eukaryotic cells to provide a platform for organizing both signal transduction^12–15^ and signal transmission^10,11,16,17^. The protein composition of primary cilia is dynamically mediated by intraflagellar transport (IFT), which employs microtubule-binding motor proteins and cargo adaptor proteins to translocate signaling components along the ciliary ultrastructure^18–22^. Certain IFT components play a role in ciliary EV shedding and release^9,11,23–25^, although the specific contribution of individual IFT motor proteins to selective cargo enrichment has not been explored.

In *Caenorhabditis elegans*, EVs are shed from primary cilia of select sensory neurons, including the inner labial type 2 (IL2) and amphid sensory neurons, as well as the male-specific ray type B (RnB), hook B (HOB), and cephalic male (CEM) neurons^11,26^. EVs shed from the ciliary distal tip of male-specific sensory neurons are released through a cuticular pore into the environment and enter the hermaphrodite uterus during copulation^23,27^ and can modulate animal behavior^11^. Conversely, EVs shed from the periciliary membrane compartment (PCMC) at the ciliary base can be phagocytosed by adjacent glial cells to help maintain cilia structure and function^23,26^. Very few ciliary EV cargo proteins have been examined in detail^28^, limiting our understanding of mechanisms underlying cargo enrichment into these EVs.

Here, we identified the ion channel CLHM-1 in EVs released from both male and hermaphrodite ciliated sensory neurons. In the absence of mating partners, release of CLHM-1-containing EVs from males decreased, suggesting that this process is a physiological response to mate availability. Visualizing EVs containing different fluorescently labeled cargoes, we discovered that discrete ciliary localization of CLHM-1 resulted in enrichment in an environmentally released EV subpopulation distinct from previously described EVs. Loss of individual kinesin-II motors had differential impact on protein colocalization in cilia and EVs. Unlike other known EV cargoes, complete elimination of kinesin-II motor protein activity did not inhibit CLHM-1 cilia entry or inclusion in EVs. Our results highlight CLHM-1 as a unique *C. elegans* ciliary EV cargo and demonstrate that anterograde IFT is a selective mechanism for cargo enrichment.

## RESULTS

### CLHM-1 is cargo in EVs released from *C. elegans* sensory neuron cilia

*<u>Cal</u>cium <u>h</u>omeostasis <u>m</u>odulator 1* (*CALHM1*) is an ion channel that modulates neuron excitability^29,30^, and human genetic studies suggest that the P86L polymorphism in *CALHM1* lowers the age of Alzheimer’s disease onset^31,32^. CALHM1 and its sole *C. elegans* homolog, CLHM-1, exhibit similar biophysical properties and functional conservation^33,34^. *clhm-1* was found to be expressed in a subset of ciliated neurons, including the IL2, ASE, ASG, ASI, ASJ, ASK, ADE, PHA, and PHB neurons (Fig. 1A)^33^. To determine if *clhm-1* is also expressed in male-specific neurons, we created transgenic animals coexpressing *gfp* from the *clhm-1* promoter and *mCherry* from the *klp-6* promoter, which drives expression in IL2 and male ciliated sensory neurons^35^. Colocalization analysis showed that *clhm-1* is expressed in the CEM neurons in the male head and HOB and RnB neurons in the male tail (Fig. 1A), all of which shed EVs (Fig. 1B)^11,23,24,28,36^.

**Fig. 1.**
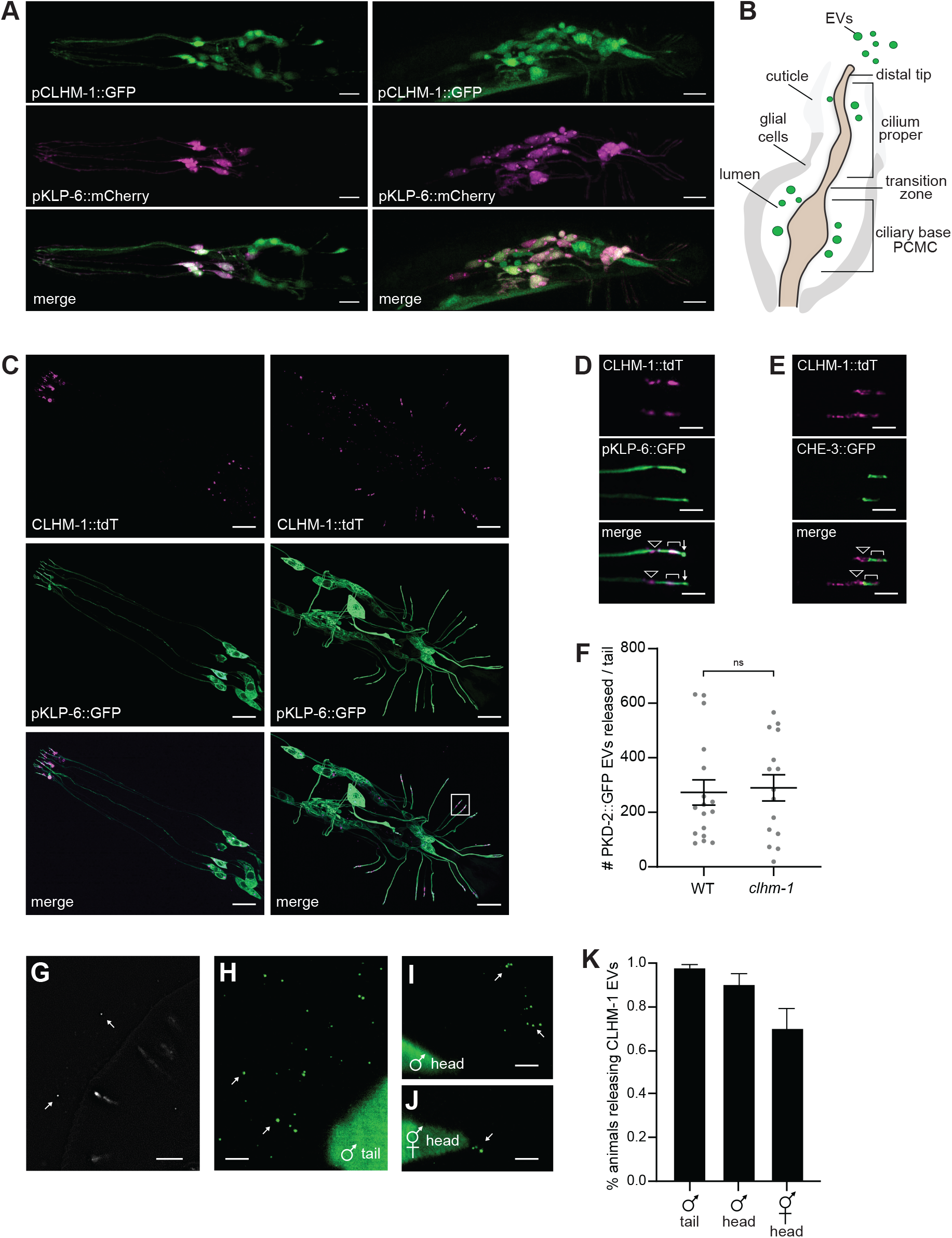
CLHM-1 is cargo in EVs released from male and hermaphrodite ciliated neurons. A) A *clhm-1* promoter::*gfp* transgene (top) drives GFP expression in male head (left) and tail (right) neurons. Colocalization with a *klp-6* promoter::*mCherry* reporter (middle) shows that *clhm-1* is expressed in IL2, CEM, RnB, and HOB EVNs (merge, bottom) plus additional sensory neurons. Scale bars, 10 µm. B) Schematic of EVs released from a *C. elegans* cilium into the environment. C) CLHM-1::tdTomato (top) localizes to neuronal cilia in the adult male head (left) and tail (right). EVNs filled out with GFP expressed from the *klp-6* promoter (middle). Bottom, merge; scale bars, 10 µm. D) CLHM-1 localizes to the ciliary base (triangle) and cilium proper (bracket), but is excluded from the distal tip (arrow); RnB4 and RnB5 (box, panel C); scale bars, 3 µm. E) CLHM-1::tdTomato (top) colocalizes with CHE-3::GFP (middle) in RnB4 and RnB5 cilia proper. Labels and scale as in (D). F) Average number of PKD-2::GFP EVs released from the male tail does not change between wild type and *clhm-1(tm4071)* animals. Error bars show SEM, Mann-Whitney test; n ≥ 15. Structured illumination microscopy shows CLHM-1::GFP localized to RnB cilia and released in EVs (arrows). Scale bar, 5 µm. H-J) TIRF microscopy shows CLHM-1::GFP EVs (arrows) released from the male tail (H), male head (I), and hermaphrodite head (J). Scale bars, 3 µm. K) Percent of males and hermaphrodites releasing CLHM-1 EVs. n ≥ 23.

To define CLHM-1 neuronal localization, we coexpressed tdTomato-tagged CLHM-1 with a *klp-6* promoter::*gfp* transcriptional reporter and observed CLHM-1 in the PCMC of the ciliary base and middle segment of the cilium proper (Fig. 1C-D). We confirmed this localization pattern by coexpression of CLHM-1::tdTomato with GFP-tagged CHE-3 (Fig. 1E), a dynein-2 heavy chain which localizes to non-motile cilia^37^. Based on the localization of CLHM-1 to cilia of EV-releasing neurons (EVNs) and the permeability of CALHM channels to Ca^2+^ and ATP^29,33,38,39^, both of which play a role in EV shedding^40–42^, we hypothesized that CLHM-1 could regulate EV biogenesis. To test this, we crossed GFP-tagged PKD-2, a conserved TRP polycystin-2 channel and well-characterized cargo present in cilia tip-derived EVs^11,23,27^, with *clhm-1(tm4071)*. Loss of *clhm-1* had no effect on the number of PKD-2 containing EVs released into the environment (Fig. 1F). This suggests that CLHM-1 is not required for release of these ciliary EVs under basal conditions.

We next considered that CLHM-1 could be packaged into ciliary EVs. Using structured illumination and total internal reflection fluorescence (TIRF) super-resolution microscopy, we imaged animals expressing CLHM-1::GFP at single copy and discovered that CLHM-1 is cargo in EVs released from male tail EVNs (Fig. 1G,H). Using lambda spectral imaging followed by linear unmixing, we confirmed the signal from EVs as bona fide GFP emission (Fig. S1). CLHM-1::GFP EV release was not restricted to males, but also observed from hermaphrodites (Fig. 1I-K). Since CLHM-1 tagged with GFP is functional and expressed at endogenous levels^33^, this localization to EVs is not the result of aberrant protein folding or over-expression. Further, not all ciliary-localized proteins in these neurons are found in EVs^11,23,36^, indicating that CLHM-1 is selectively packaged.

### CLHM-1 and PKD-2 cargoes are enriched in distinct EV subpopulations

Expression of PKD-2 is restricted to male-specific neurons, while CLHM-1 is expressed in a wider subset of both male and hermaphrodite EVNs^33,43^. While we observed release of CLHM-1::GFP EVs from nearly every animal of both sexes, fewer EVs were detected in images of males expressing CLHM-1::GFP compared to PKD-2::GFP. As a single cilium can release multiple EV subpopulations^23,28^, we hypothesized that CLHM-1 could be enriched in a different environmentally-released EV subset than PKD-2. We created animals expressing single-copy insertion (SCI) fluorescent transgenes in a <u>h</u>igh incidence of <u>m</u>ales *him-5(e1490)* background^44^: one strain coexpresses CLHM-1::GFP with CLHM-1::tdTomato, a second strain coexpresses PKD-2::GFP with PKD-2::tdTomato, and two strains coexpress PKD-2::GFP with CLHM-1::tdTomato. Using TIRF super-resolution microscopy, we imaged EVs released from male tail EVNs, as these EVs are abundant and accumulate in the hermaphrodite uterus during mating, indicating physiological potential^27,28^. Transgene expression was driven by appropriate promoters to ensure overlap in the tail EVNs. CLHM-1::GFP largely colocalized with CLHM-1::tdTomato in environmentally-released EVs (Fig. 2A), as did PKD-2::GFP and PKD-2::tdTomato (Fig. 2B). In contrast, CLHM-1::tdTomato and PKD-2::GFP were observed together in fewer EVs (Fig. 2C).

**Fig. 2.**
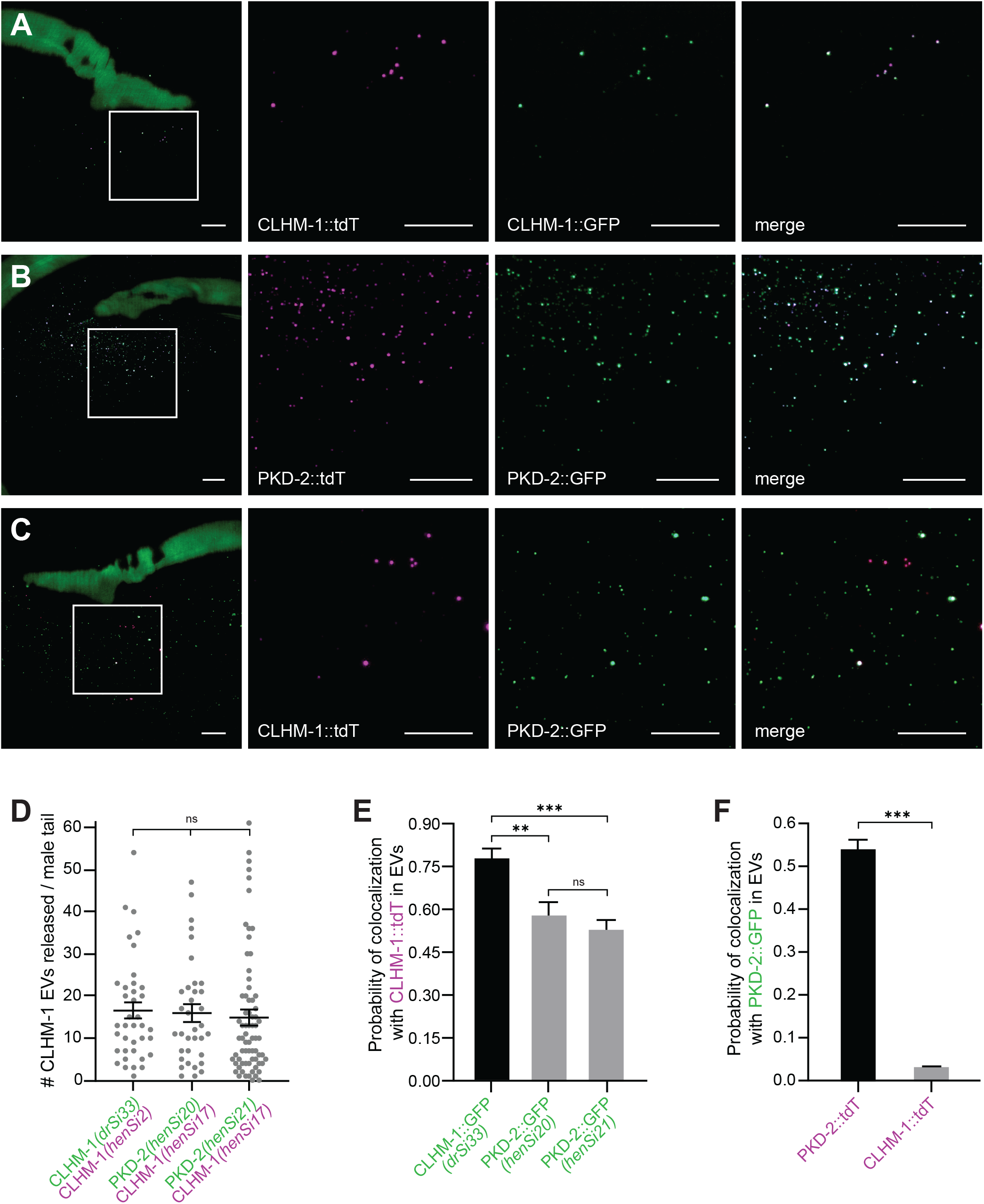
CLHM-1 and PKD-2 are enriched in distinct EV subpopulations released from male tail EVNs. A) CLHM-1::tdTomato (tdT; *henSi2*) and CLHM-1::GFP (*drSi33*) colocalize in EVs released into the environment. Boxed region (left) is enlarged in subsequent images to clearly show EVs. B) PKD-2::tdTomato (*henSi26*) and PKD-2::GFP (*henSi20*) colocalize in EVs. C) PKD-2::GFP (*henSi20*) and CLHM-1::tdTomato (*henSi17*) display less frequent colocalization in EVs (compare to A,B). D) Average number of CLHM-1::tdTomato EVs released is not different between animals expressing different CLHM-1::tdTomato transgenes (*henSi2, henSi17*) or when CLHM-1::tdTomato is paired with different PKD-2::GFP SCI transgenes (*henSi20, henSi21*); n ≥ 35. E) Probability of GFP cargo colocalization with CLHM-1::tdTomato in EVs. CLHM-1::GFP is more likely than PKD-2::GFP to coincide with CLHM-1::tdTomato; n ≥ 35. F) Probability of tdTomato cargo colocalization with PKD-2::GFP in EVs. PKD-2::tdTomato is more likely than CLHM-1::tdTomato to coincide with PKD-2::GFP; n ≥ 29. Scale bars, 10 µm. Kruskal-Wallis test used for (D,E). Mann-Whitney test used for (F). Error bars show SEM; ** = p<0.01, *** = p<0.001.

Using the Imaris spot detector, calibrated to minimize noise detection, we identified EVs with GFP or tdTomato signal, as well as EVs with signal overlap (Fig. S2). The number of environmentally-released EVs containing CLHM-1 was not different between strains carrying *henSi2* or *henSi17*, two different SCI CLHM-1::tdTomato transgenes (Fig. 2D). Additionally, pairing a CLHM-1::tdTomato transgene with a CLHM-1::GFP SCI transgene (*drSi33*) or two different PKD-2::GFP SCI transgenes (*henSi20* and *henSi21*) did not alter the number of CLHM-1::tdTomato EVs detected (Fig. 2D). This shows that the presence of GFP-labeled CLHM-1 does not alter the number of detectable CLHM-1::tdTomato EVs and demonstrates that our imaging and analysis pipeline generates reproducible quantitative data with multiple independent transgenes.

We next examined signal overlap in EVs and discovered that CLHM-1::GFP was significantly more likely than PKD-2::GFP to colocalize with CLHM-1::tdTomato, despite PKD-2::GFP being the more abundant EV cargo (Fig. 2E). Likewise, PKD-2::tdTomato was significantly more likely than CLHM-1::tdTomato to colocalize with PKD-2::GFP in EVs (Fig. 2F). These data indicate that PKD-2 and CLHM-1 are deliberately enriched into distinct subpopulations of ciliary EVs shed from the same sensory neuron cilia.

### Mating partner presence differentially impacts release of PKD-2 and CLHM-1 containing EVs

Given the observation of CLHM-1 and PKD-2 in distinct EV subpopulations, we wondered if these EVs were differentially shed in response to environmental cues. The presence of hermaphrodite mating partners has been shown to modulate relative abundance of PKD-2-containing EVs^23^. We compared the number of CLHM-1::tdTomato-containing EVs shed from tail sensory neurons of males cultured with adult hermaphrodites to the number released by virgin males separated from hermaphrodites at the fourth larval (L4) stage. The absence of mating partners significantly reduced CLHM-1 EV release (Fig. 3A). Conversely, release of PKD-2 EVs from virgin males was significantly higher compared to males cultured with hermaphrodites (Fig. 3B, Fig. S3). We observed a significant increase in the probability of PKD-2::GFP being present in CLHM-1::tdTomato EVs released from virgin adult males, possibly due to the concurrent increase in PKD-2 and decrease in CLHM-1 EV shedding (Fig. 3C).

**Fig. 3.**
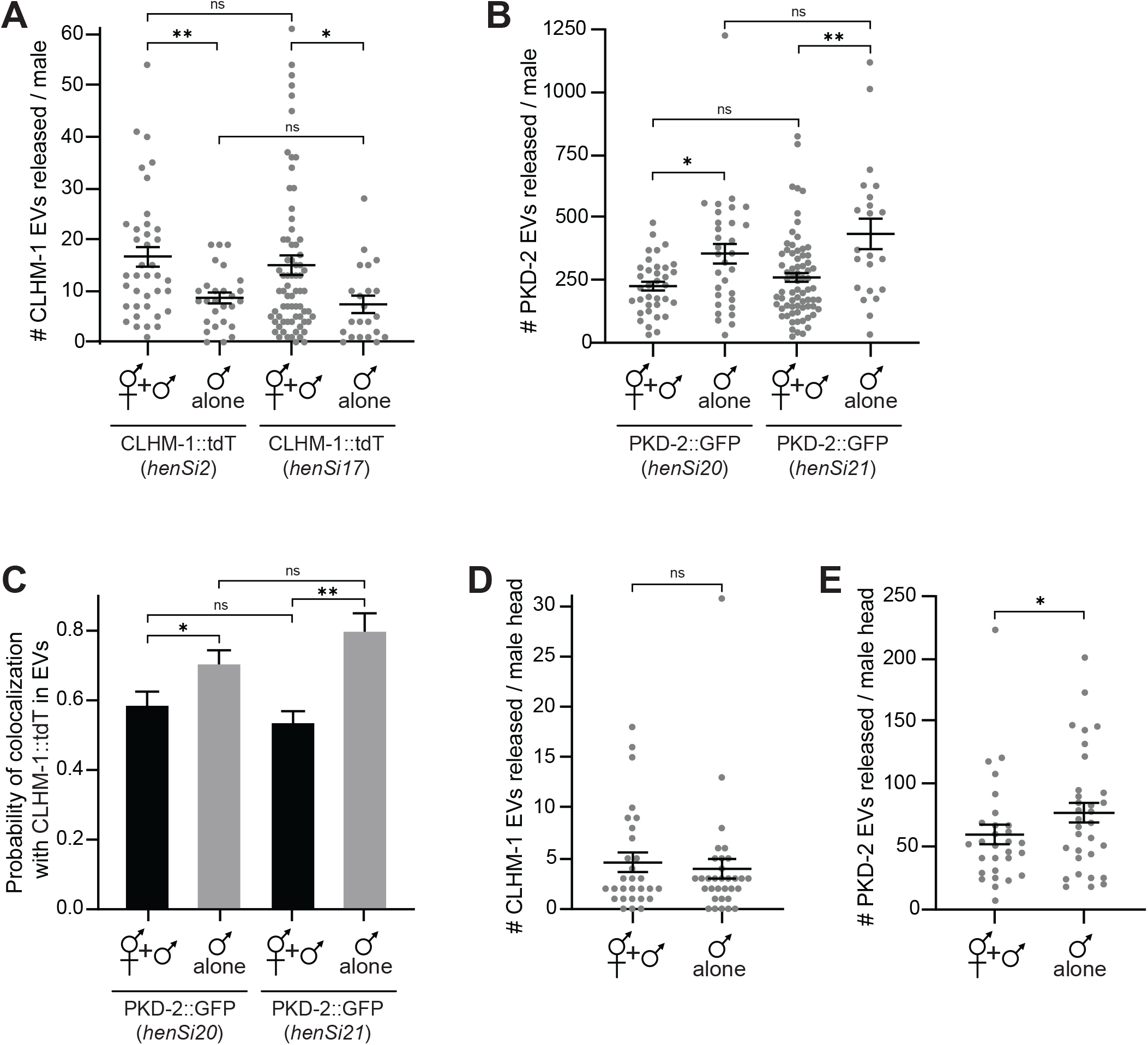
Abundance of CLHM-1 and PKD-2 containing EVs is dependent on mating partner availability. A) Average number of CLHM-1::tdTomato EVs released per virgin male tail is lower compared to males raised with mating partners, consistent between transgenes (*henSi2, henSi17*); n ≥ 22. B) Average number of PKD-2::GFP EVs released per virgin male tail is higher compared to males raised with mating partners, consistent between transgenes (*henSi20, henSi21*); n ≥ 22. C) Probability of PKD-2::GFP being present in a CLHM-1::tdTomato-containing EV is greater in EVs released from virgin male tails compared to males raised with mating partners; n ≥ 20. D) Availability of mating partners does not affect release of CLHM-1::tdTomato (*henSi17*) EVs from the male head; n ≥ 29. E) Average number of PKD-2::GFP (*henSi21*) EVs released per virgin adult male head is higher compared to males raised with mating partners; n ≥ 29. Error bars show SEM; Mann-Whitney test * = p<0.05, ** = p<0.01.

Male tail neurons respond to mechanosensory cues, while many head EVNs are chemosensory^45–47^. Thus, we sought to determine if mating partner availability had differential impact on EV release from neurons that respond to different stimuli. Hermaphrodite presence did not impact release of CLHM-1 EVs from male head EVNs (Fig. 3D). However, just as observed for the tail neurons, PKD-2 EV release from head EVNs of virgin males was significantly higher than from males cultured with hermaphrodites (Fig. 3E, Fig. S3). In conclusion, CLHM-1 and PKD-2 EV release is differentially altered by mating partner availability, demonstrating that cargo abundance within these EVs is a dynamically regulated and not just the result of non-specific ciliary membrane shedding.

### PKD-2, but not CLHM-1, localizes to the distal segment of male tail EVN cilia

Primary cilia contain multiple distinct compartments, each with unique ultrastructure, protein enrichment and lipid composition. In *C. elegans* EVNs, the distal dendrite gives way to a specialized PCMC at the ciliary base and the transition zone separates the ciliary base from the cilium proper^48–52^. While both PKD-2 and CLHM-1 are ciliary proteins^33,43^, we reasoned that enrichment of these cargoes in distinct EV subpopulations could result from differential localization to specific ciliary compartments or microdomains.

To define relative CLHM-1 and PKD-2 localization, we imaged RnB neurons in males coexpressing CLHM-1::tdTomato with PKD-2::GFP. While both CLHM-1 and PKD-2 localized to the PCMC and middle segment of the cilium proper, PKD-2::GFP alone extended into the distal tip (Fig. 4A,B). We quantified colocalization between CLHM-1::tdTomato and either CLHM-1::GFP or PKD-2::GFP in three-dimensional RnB cilia reconstructions. The Manders 1 (M1) colocalization coefficient, a measure of green pixels that co-occur with red pixels, was higher in the cilium proper when comparing overlap of CLHM-1::GFP with CLHM-1::tdTomato to overlap of PKD-2::GFP with CLHM-1::tdTomato (Fig. 4C). We also analyzed PKD-2::GFP and CLHM-1::tdTomato fluorescence intensity along the cilia in two-dimensional image projections. Distribution profiles showed that PKD-2::GFP extended into the distal tip whereas CLHM-1::tdTomato was absent, consistent with our visual observations and three-dimensional analyses (Fig. 4D). We frequently observed EVs containing only PKD-2 being shed from the distal tip (Fig. 4E) as previously described^23^. Thus, the absence of CLHM-1 from this ciliary subcompartment provides a mechanism for enrichment of PKD-2 cargo into one EV subpopulation.

**Fig. 4.**
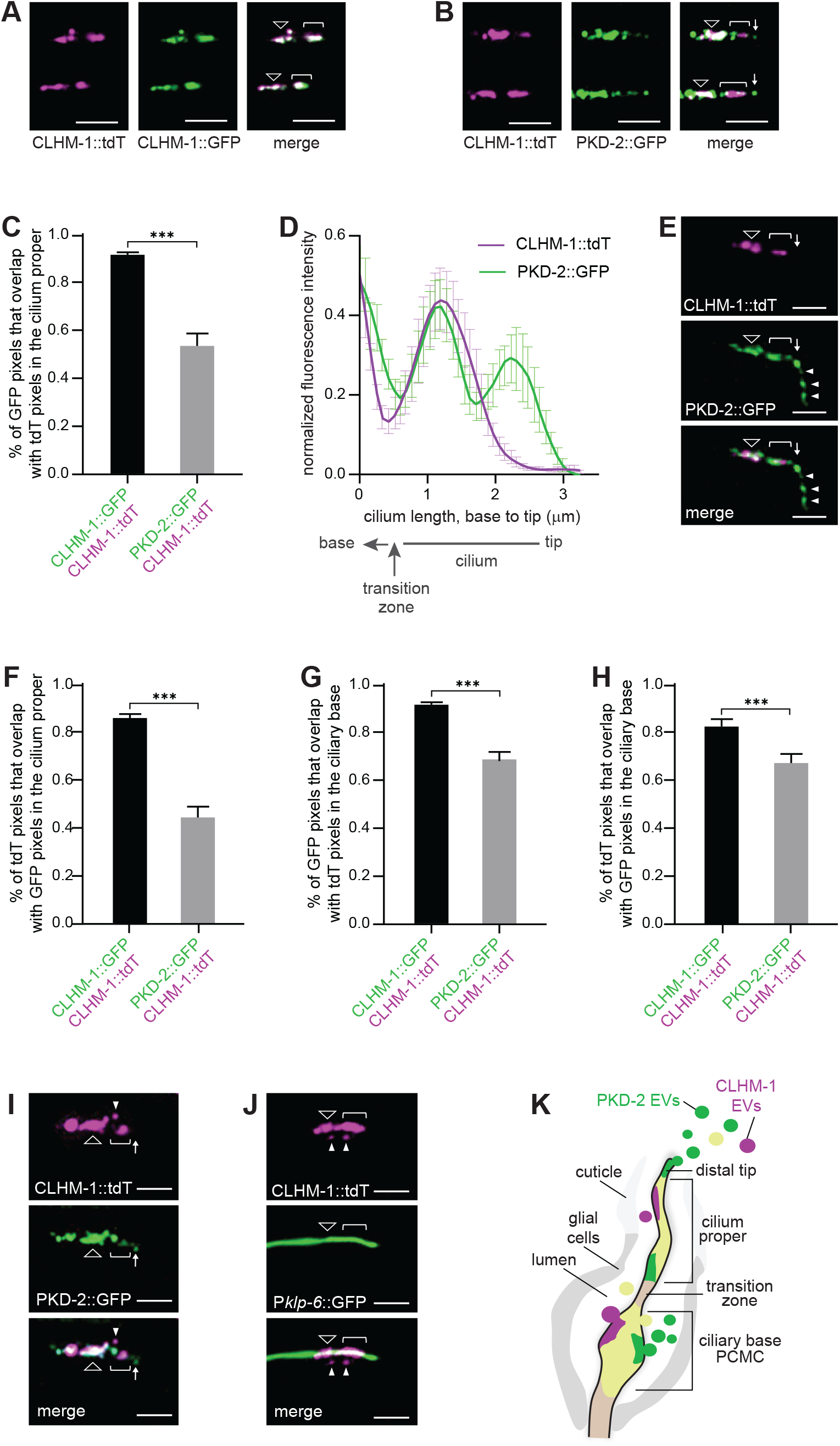
EVs released into the environment are shed from multiple ciliary subcompartments. A) CLHM-1::tdTomato (left) and CLHM-1::GFP (middle) localize to the ciliary base (triangle) and the middle segment (bracket). Representative RnB4 and RnB5 cilia shown; scale bars, 3 µm. B) CLHM-1::tdTomato (left) and PKD-2::GFP (middle) colocalize in the ciliary base (triangle) and middle segment (bracket). PKD-2::GFP localizes to the distal tip (arrow) of RnB4 and RnB5 cilia; scale bars, 3 µm. C) In the cilium proper, the M1 colocalization coefficient is higher when CLHM-1::tdTomato is expressed with CLHM-1::GFP rather than PKD-2::GFP; n ≥ 25. D) Normalized fluorescence intensity of PKD-2::GFP and CLHM-1::tdTomato along RnB cilia, measured from base to tip (left to right) shows PKD-2, but not CLHM-1, enters the distal tip; n = 16. E) PKD-2::GFP (middle), but not CLHM-1::tdTomato (top) is present in EVs (arrowheads) shed from the distal tip (arrow). Scale bars, 2 µm. F) The M2 colocalization coefficient in the cilium proper is higher when CLHM-1::tdTomato is expressed with CLHM-1::GFP rather than PKD-2::GFP; n ≥ 25. G-H) M1 (G) and M2 (H) are higher in the ciliary base when CLHM-1::tdTomato is expressed with CLHM-1::GFP compared to PKD-2::GFP; n ≥ 25. I) An EV (arrowhead) containing CLHM-1::tdTomato (top), but not PKD-2::GFP (middle) is shed from the PCMC of the ciliary base (triangle) in relation to the cilium proper (bracket) and distal tip (arrow); scale bars, 2 µm. J) EVs (arrowheads) containing CLHM-1::tdTomato (top) are shed from the PCMC of the ciliary base (triangle) and observed adjacent to the middle segment of the cilium proper (bracket). EVNs are filled out with GFP expressed from the *klp-6* promoter (middle); scale bars, 2 µm. K) Schematic of discrete ciliary membrane regions shedding EVs containing only CLHM-1 (magenta), only PKD-2 (green), or both (yellow) cargoes. Error bars show SEM; Mann-Whitney test, *** = p<0.001.

We observed regions in the cilium middle segment and PCMC that contained only tdTomato signal (Fig. 4B) that cannot be described by the M1 coefficient, which only provides colocalization information for pixels containing GFP. The Manders 2 (M2) coefficient, which indicates the percentage of red pixels that co-occur with green pixels, was lower in the cilium proper of animals that expressed CLHM-1::tdTomato with PKD-2::GFP compared to CLHM-1::GFP, suggesting that regions of the cilium proper contain CLHM-1 but not PKD-2 (Fig. 4F). Both M1 and M2 colocalization coefficients were also lower in the PCMC when CLHM-1::tdTomato was coexpressed with PKD-2::GFP compared to CLHM-1::GFP (Fig. 4G,H). These data suggest that CLHM-1 is enriched in discrete membrane domains of multiple different ciliary compartments.

The PCMC at the ciliary base of *C. elegans* sensory neurons is a site of EV shedding, however, these EVs were previously not thought to be released into the environment^23,24,26^. Since our data show that CLHM-1-containing EVs are not tip-derived yet are still released into the environment, these EVs must originate from a secondary site(s). We often observed shedding of CLHM-1 EVs from the PCMC (Fig. 4A,I,J), though it remains possible that CLHM-1 EVs could also bud from the cilium middle segment. In conclusion, we suggest that enrichment of CLHM-1 and PKD-2 into discrete subpopulations of EVs derived from multiple sites on the same cilium is achieved by differential ciliary localization (Fig. 4K).

### Kinesin-II motors specify EV cargo enrichment and biogenesis

IFT, the bi-directional movement of proteins along ciliary microtubules, is required for establishing and maintaining cilium ultrastructure and compartmentalization^20^. Anterograde IFT in *C. elegans* cilia is driven by homodimeric kinesin OSM-3 and a heterotrimeric kinesin comprised of KLP-11, KLP20, and KAP-1^53,54^. In IL2 and male-specific EVNs, the kinesin-3 KLP-6 also participates in anterograde IFT^55^. As IFT is required for translocation of many ciliary membrane proteins, we considered that this could serve as a mechanism to enrich PKD-2 and CLMH-1 into different ciliary regions and thus, EV subpopulations.

We found that the average number of PKD-2::GFP EVs released from male tail EVNs did not change between wild type, *osm-3(mn391), osm-3(p802)*, and *klp-11(tm324)* mutants (Fig. 5A). Analysis of ciliary PKD-2::GFP distribution showed that PKD-2 was still transported to the RnB distal tip in both *klp-11* and *osm-3* mutants (Fig. 5B,C), consistent with lack of impact of these mutations on EV release. Surprisingly, loss of either *osm-3* or *klp-11* significantly increased release of EVs containing CLHM-1 (Fig. 5D) without impacting CLHM-1::tdTomato ciliary distribution (Fig. 5B,C). Loss of either *osm-3* or *klp-11* did not affect average CLHM-1::tdTomato intensity in the cilium proper, but did cause a small decrease in fluorescence intensity in the PCMC (Fig. 5E,F). This suggests that the kinesin-dependent increase in CLHM-1-containing EV shedding is not a result of ciliary mislocalization.

**Fig. 5.**
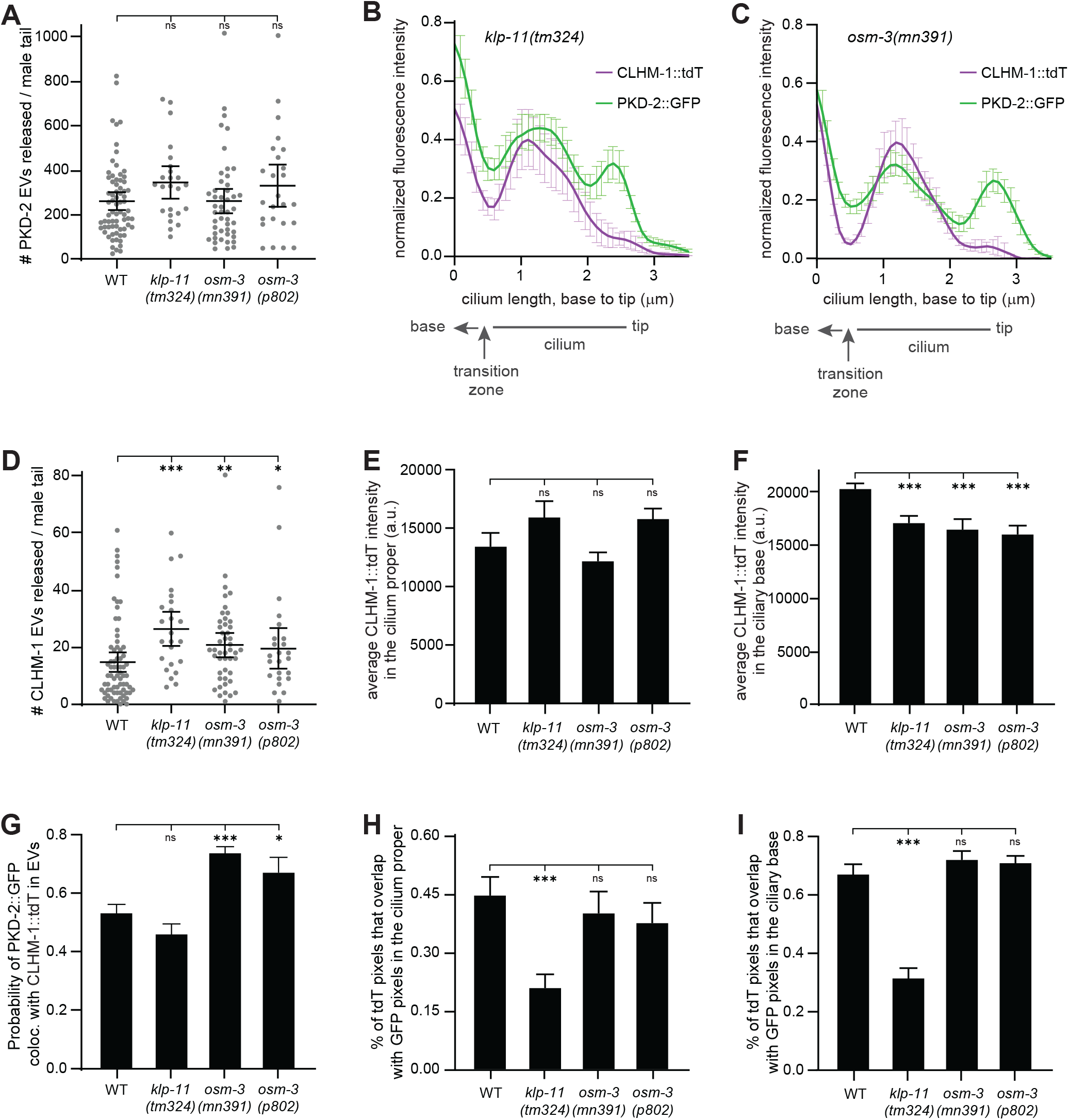
Loss of OSM-3, but not KLP-11, affects enrichment of ciliary EV cargo. A) Average number of PKD-2::GFP EVs released from male tail EVNs does not change in *klp-11* or *osm-3* mutants compared to wild type; n ≥ 24. B-C) Normalized fluorescence intensity of PKD-2::GFP and CLHM-1::tdTomato along RnB cilia in *klp-11* (B) or *osm-3* (C) mutants. PKD-2, but not CLHM-1, enters the distal tip as in wild type; n = 15. D) Average number of released CLHM-1::tdTomato EVs increases in *klp-11* and *osm-3* mutants compared to wild type; n ≥ 24. E) CLHM-1::tdTomato fluorescence intensity in the cilium proper of *klp-11* and *osm-3* mutants is the same as wild type; n ≥ 16. F) CLHM-1::tdTomato fluorescence intensity in the ciliary base is lower in *klp-11* and *osm-3* mutants compared to wild type; n ≥ 16. G) Probability of PKD-2::GFP being present in a CLHM-1::tdTomato-containing EV does not change in *klp-11* mutants but increases in *osm-3* mutants; n ≥ 24. H,I) The M2 coefficient significantly decreases in the cilium proper (I) and ciliary base (J) of *klp-11* mutant males, indicating reduced colocalization of CLHM-1::TdT with PKD-2::GFP. M2 was unchanged in *osm-3* mutants compared to wild type; n ≥ 21. Error bars show SEM; Kruskal-Wallis test * = p<0.05, ** = p<0.01, *** = p<0.005.

We next examined colocalization of CLHM-1 and PKD-2 in EVs released from the IFT mutants. The probability of PKD-2::GFP being found in a CLHM-1::tdTomato EV increased in both *osm-3* mutants but was unchanged in the *klp-11* mutant compared to wild type (Fig. 5G). While trafficking of CLHM-1 and PKD-2 to ciliary compartments was not affected by loss of KLP-11 or OSM-3 (Fig. 5B,C), we sought to determine if these kinesins impacted CLHM-1::tdTomato and PKD-2::GFP colocalization within different ciliary regions. Compared to wild type, we observed a significant decrease in the M2 coefficient in both the cilium proper and PCMC of *klp-11* mutants, indicating an increase in CLHM-1::tdTomato that did not colocalize with PKD-2::GFP (Fig. 5H,I). No change in ciliary colocalization of CLHM-1 and PKD-2 was detected in either *osm-3* mutant (Fig.5H,I). These data show an interesting correlation between the ciliary colocalization of CLHM-1 and PKD-2 and their coincidence in EVs. The increase in CLHM-1 EV release in the *osm-3* mutant is associated with an increase in colocalization of PKD-2 and CLHM-1 in EVs. Conversely, loss of *klp-11* reduces ciliary colocalization of these cargoes, suggesting that the additional CLHM-1-containing EVs released in the *klp-11* mutant are shed from membrane regions that contain only CLHM-1.

*C. elegans* male-specific EVNs also employ the kinesin-3 KLP-6^55,56^, which is important for release of PKD-2 EVs into the environment, but does not prevent ciliary EV shedding into the lumen^11^. We observed a significant decrease in CLHM-1 EVs released into the environment in the *klp-6* mutant compared to the wild type (Fig. S4), suggesting that KLP-6 is required for environmental release for multiple EV subpopulations. Together, our data demonstrate that each kinesin plays a distinct role, differentially impacting ciliary colocalization and enrichment of EV cargoes as well as release of EV subpopulations into the environment.

### Anterograde IFT is required for release of EVs containing PKD-2, but not CLHM-1

Loss of anterograde IFT in *osm-3*; *klp-11* double mutants dramatically decreases shedding of EVs containing PKD-2::GFP^11,23^ without impacting the length of male-specific CEM cilia^55^. To determine if release of other ciliary EV subpopulations is reliant on the cooperative action of these kinesin motors, we imaged PKD-2::GFP and CLHM-1::tdTomato EVs shed from wild type and *osm-3*; *klp-11* double mutants (Fig. 6A,B). Quantification showed a severe reduction in release of PKD-2-containing EVs in the double mutant as previously reported^11,23^, while the number of CLHM-1 EVs released into the environment did not significantly change (Fig. 6C,D). We also observed a significant decrease in the probability of PKD-2::GFP being present with CLHM-1::tdTomato in EVs released from *osm-3*; *klp-11* animals (Fig. 6E). This shows that loss of the cooperating OSM-3 and heterotrimeric kinesin-II motors has a subpopulation-specific impact on EV release.

**Fig. 6.**
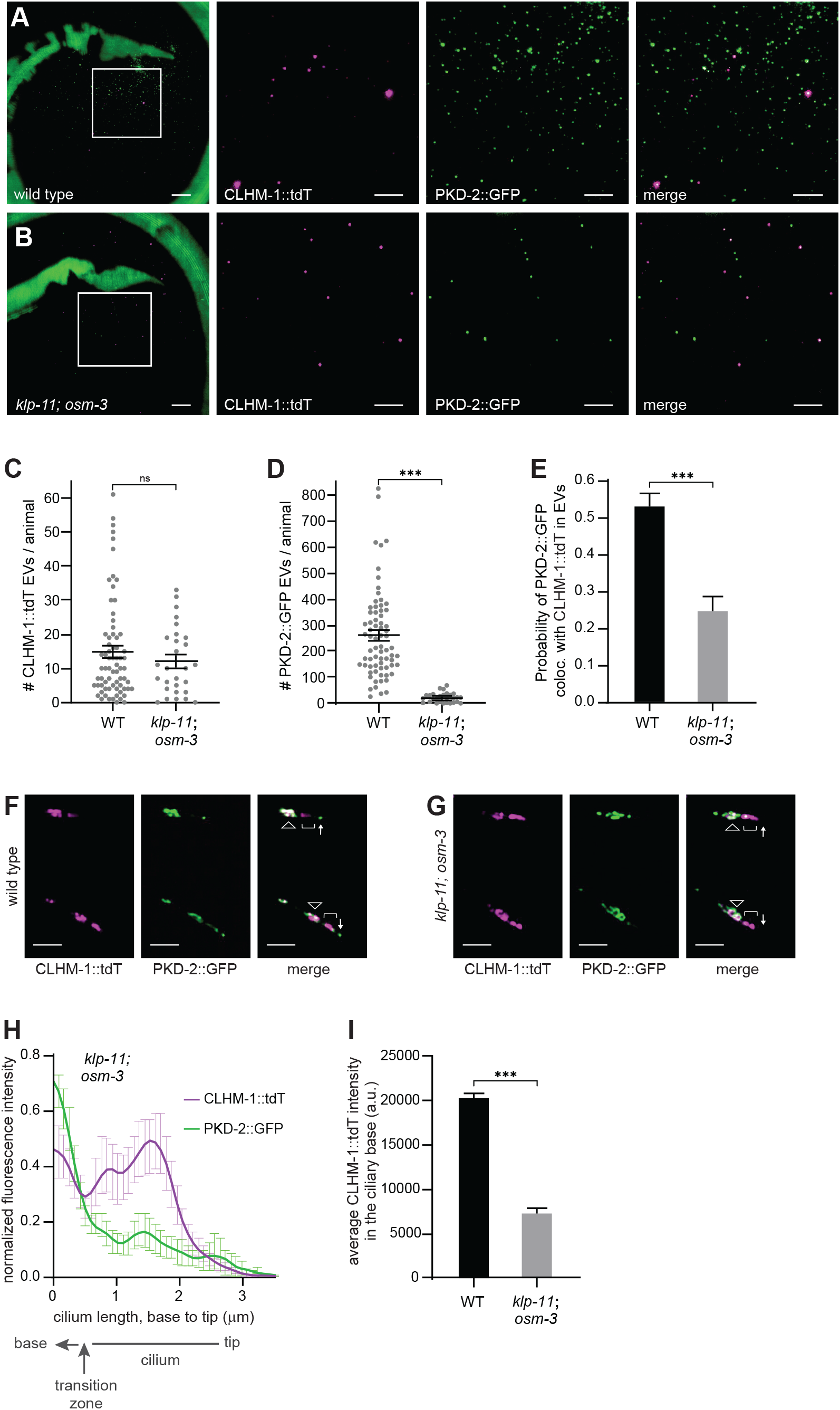
Loss of kinesin-II activity reduces release of PKD-2, but not CLHM-1 EVs. A-B) Representative images of CLHM-1::tdTomato (*henSi17*) and PKD-2::GFP (*henSi21*) EVs released from wild type (A) and *klp-11; osm-3* (B) male tails. Boxed region (left) is enlarged in subsequent images; scale bars, 3 µm. C) CLHM-1::tdTomato EV release is not altered in *klp-11; osm-3* mutants; n ≥ 28. D) Release PKD-2::GFP EVs decreases in *klp-11; osm-3* mutants; n ≥ 28. E) Probability of PKD-2::GFP presence in CLHM-1::tdTomato EVs decreases in *klp-11; osm-3* mutants; n ≥ 28. F,G) PKD-2::GFP localizes to the distal tip (arrow) in wild type, but not *klp-11*; *osm-3* mutants. CLHM-1::tdTomato localizes to the PCMC (triangle) and cilium proper (bracket) in both wild type and *klp-11; osm-3* animals. Representative RnB2 and RnB3 cilia shown; scale bars, 3 µm. H) Normalized fluorescence intensity of PKD-2::GFP and CLHM-1::tdTomato along RnB cilia in *klp-11*; *osm-3* mutants; compare to wild type (Fig 4E); n = 14. I) CLHM-1::tdTomato fluorescence intensity in the ciliary base is significantly reduced in the *klp-11*; *osm-3* mutant compared to wild type; n ≥ 16. Error bars show SEM; Mann-Whitney test, *** = p<0.001.

We imaged RnB cilia in wild type and *osm-3*; *klp-11* animals to determine if changes in protein distribution could shed light on the changes in EV release and cargo colocalization. While CLHM-1::tdTomato was still trafficked into the cilium proper, PKD-2::GFP was often excluded entirely and rarely found at the distal tip in *osm-3; klp-11* mutants (Fig. 6F,G). This observation, confirmed by quantification of fluorescence intensity distribution profiles along cilia (Fig. 6H), suggests that shedding of PKD-2-containing EVs from the distal tip is largely abolished in the double mutant. CLHM-1::tdTomato fluorescence intensity in the cilium proper was unchanged between wild type and *klp-11; osm-3* mutants, showing that cilia entry of CLHM-1 occurs independently of these kinesin motors (Fig. S5). However, a decrease in CLHM-1::tdTomato fluorescence intensity was observed in the PCMC, suggesting that loss of OSM-3 and heterotrimeric kinesin-II have an additive impact on CLHM-1 abundance in this compartment (Fig. 6I). Our data suggest that discrete ciliary localization, mediated by IFT, results in specific protein enrichment into EVs shed from primary cilia.

## DISCUSSION

Cilia protruding from cells act not only to receive signals, but also to transmit signals via EV release. It is important to understand how proteins become enriched in or excluded from specific ciliary EV subpopulations as the physiological potential of an EV is derived from its cargo^1,7,10^. Using *C. elegans*, we identified the ion channel CLHM-1 as a new cargo in ciliary-derived EVs released from sensory neuron cilia into the environment. While EVs are also shed from the distal tip of primary cilia^8,9,23^, CLHM-1 was excluded from this ciliary compartment and thus, tip-derived EVs. The presence of mating partners caused an increase in tip-derived EV release, but a significant decrease in shedding of EVs containing CLHM-1. This indicates that localization of cargoes to ciliary subcompartments allows for heterogeneous EV subpopulations to be dynamically and differentially shed in response to physiological stimuli.

Since there are multiple different ciliary-derived EVs subpopulations, this raises the question of how proteins are discretely localized to ensure enrichment into distinct subsets. Heterotrimeric kinesin-II and homomeric OSM-3 move IFT trains from the ciliary base, through the transition zone and up to the distal tip^57,58^. In the absence of these kinesin motors, many ciliary proteins accumulate at the ciliary base ^23,55^, though not all proteins are dependent on kinesin-II-mediated IFT for entry into the cilium proper^59,61^. We and others found that PKD-2 fails to localize to the distal tip and is not released in tip-derived EVs in *osm-3; klp-11* mutants^23^. The myristolated protein CIL-7, another EV cargo^60^, does not require *osm-3* and *klp-11* for transport into the cilium proper, but does depend on these IFT motors for inclusion in tip-derived EVs^23^. Neither CLHM-1 abundance in the cilium proper or EV release was altered in the *osm-3; klp-11* mutant. This is consistent with CLHM-1 being excluded from tip-derived EVs and shows that anterograde IFT is required for EV shedding from the cilium distal tip, but not secondary sites. Remarkably, we revealed differential roles for individual kinesin motors in regulating ciliary EV cargo enrichment, as loss of *osm-3* versus *klp-11* differentially affected colocalization of CLHM-1 and PKD-2 in the cilium and released EVs. We propose that activity of specific kinesin motors differentially impacts the localization of still unknown proteins required for cargo enrichment and EV shedding.

It has been suggested that packaging of proteins into EVs serves as a mechanism to reduce accumulation of cargoes resulting from disrupted IFT or protein overexpression^26^. However, our results are not consistent with all ciliary EVs being used only as vectors for disposal. First, our fluorescent EV cargoes were expressed at single copy, yet we still observed frequent and abundant shedding of EVs into the environment. Second, while more PKD-2::GFP EVs were detected from animals containing an overexpression transgene, we observed the same relative change in release of overexpressed and endogenously-expressed PKD-2::GFP EVs in response to mating partner availability. Third, CLHM-1 transport into the cilium proper was not dependent on kinesin-II activity, and instead of observing greater abundance, we found decreased CLHM-1 accumulation in the ciliary base of the kinesin mutants. Finally, no increase in shedding of CLHM-1 EVs was observed in the *klp-11; osm-3* double mutant. Together, these data indicate that IFT disruption is not a main driver of ciliary EV shedding and environmental release.

Shedding of EVs containing CLHM-1 was differentially altered not only in the kinesin mutants, but also by mating partner availability, suggesting that release of these EVs into the environment is physiologically significant. Heterogeneous EVs are released from many *C. elegans* cell types^28,62^ and we currently lack a way to specifically isolate the CLHM-1-containing EV subpopulation, limiting our ability to directly interrogate the function of this protein in EVs. However, we propose two possibilities, which are not mutually exclusive. First, CALHM channels are permeable to Ca^2+^ and ATP^33,38,39^, thus CLHM-1 could act as a release channel for ions or small molecules stored in the EV lumen. Resting ciliary [Ca^2+^] is significantly higher than resting cytoplasmic [Ca^2+^]^15^, and ATP synthase subunits localize to primary cilia^63,64^, indicating that these molecules are concentrated in cilia lumen and could be captured in ciliary EVs. The second possibility posits CLHM-1 as a modulator of primary cilia signaling, where it could act to amplify signals. Excess CLHM-1 may need to be jettisoned from the cilia into EVs to prevent toxicity^33^. Regardless of the specific role for CLHM-1 in ciliary-derived EVs, our identification of this new EV subpopulation provides us with a visual tool that allows us to explore how specificity in ciliary EV biogenesis and cargo sorting is achieved.

## Supporting information

Supplemental Figures - Clupper et al

## SUPPLEMENTAL INFORMATION

Supplemental information includes five figures and can be found with this article online.

## ACKNOWLEDGEMENTS

We thank Natalia Morsci and Maureen Barr (Rutgers University; R01 DK59418) for providing PT2332, *myIs10* [Pklp-6::KLP-6::GFP +pBX1]; *him-5(e1490)*, as an unpublished gift. Additional nematode strains were provided by the *Caenorhabditis* Genetics Center, which is supported by the NIH-ORIP (P40 OD010440). Microscopy access was supported by grants from the NIH-NIGMS (P20 GM103446), NIH-ORIP (S10 OD016361), NSF (IIA-1301765), and State of Delaware. This work was supported by NIH-NIGMS T32-GM133395 (to M.C. and R.G. as part of the Chemistry Biology Interface predoctoral training program), a University of Delaware Graduate Scholars award (to M.E.), NIH-NIGMS INBRE (P20 GM103446) Pilot Project and Core Center Access Awards (to J.E.T), and NIH-NIGMS R01 GM135433 (to J.E.T.).

## AUTHOR CONTRIBUTIONS

Conceptualization, M.C. and J.E.T.; Methodology, M.C., R.G., M.E., D.T., and J.L.C.; Formal Analysis, M.C., R.G., and J.E.T.; Investigation, M.C., R.G., M.E., and J.L.C..; Writing – original draft, M.C. and J.E.T; Writing – review and editing, M.C., R.G., M.E., D.T., J.L.C, and J.E.T.; Visualization, M.C., M.E., and J.E.T; Supervision, J.E.T.; Funding Acquisition, M.C., R.G., M.E., and J.E.T.

## STAR METHODS

### RESOURCE AVAILABILITY

#### Lead Contact

Further information and reagent requests should be directed to and will be fulfilled by the lead contact, Jessica Tanis

#### Materials Availability Statement

*C. elegans* strains generated from our studies will be provided upon request. The University of Delaware Material Transfer Agreement Request Webform will be completed when research materials are transferred to an outside party.

#### Data and Code Availability

- All original data have been deposited at Mendeley Data and are publicly available as of the date of publication
- This paper does not report original code
- Any additional information required to reanalyze the data reported in this paper is available from the lead contact upon request.

## EXPERIMENTAL MODEL AND SUBJECT DETAILS

### Nematode Culture

All strains were cultured at 20°C on Nematode Growth Media (NGM) plates seeded with *E. coli* OP50^65^. *clhm-1(tm4071)* II, *klp-6(sy511)* III, *osm-3(mn391)* IV, *osm-3(p802)* IV, *klp-11(tm324)* IV, *him-5(e1490)* V, and *lin-15(n765ts)* X mutant alleles were used.

Duplex genotyping detects *clhm-1(tm4071)* and *klp-11(tm324)*; SuperSelective genotyping^66^ detects *klp-6(sy511), osm-3(mn391)*, and *osm-3(p802)*.

## METHOD DETAILS

### Transgenesis

To examine the *clhm-1* expression pattern, pJT46, a 3 kb *clhm-1* promoter::*gfp*::*unc-54* 3’ UTR construct (50 ng/µl), was injected with pENM1, a 1.58 kb *klp-6* promoter::*mCherry*::*unc-54* 3’ UTR construct (50 ng/µl), and the *lin-15* rescuing construct pL15EK (80 ng/µl) into MT8189 *lin-15(n765ts)*. Transgenic animals were created using standard germline transformation;^67^ males were generated by heat shock.

Constructs for single-copy insertion were generated using a combination of TOPO, restriction enzyme, and Gateway cloning. pCFJ910, which contains a MCS and minimal Mos1 and Neomycin/G418 resistance sequences, was used as the backbone for all constructs made for this study including: pDT285 (3 kb *clhm-1* promoter::*clhm-1*::tdTomato::let-858 3’UTR), pDT290 (1.3 kb *pkd-2* promoter::*pkd-2*::gfp::let-858 3’UTR), pDT292 (1.3 kb *pkd-2* promoter::*pkd-2*::tdTomato::let-858 3’UTR), and pDT299 (1.6 kb *klp-6* promoter::*clhm-1*::tdTomato::let-858 3’UTR). SCI transgenes were integrated into the genome using minimal Mos1 insertion^68^. Adult hermaphrodites were injected with a DNA mix containing the construct for the desired transgene (10 ng/μL), pGH8 (*rab-3* promoter::*mCherry*::*unc-54* 3’UTR;10 ng/μL), pCFJ90 (*myo-2* promoter::*mCherry*:: *unc-54* 3’UTR; 2.5 ng/μL), pCFJ104 (*myo-3* promoter::*mCherry*:: *unc-54* 3’UTR; 10 ng/μL), pCFJ601 (*eef-1A*.*1* promoter::Mos1 transposase; 50 ng/μL), and pMA122 (*hsp-16*.*41* promoter::*peel-1*::*tbb-2* 3’UTR; 10 ng/μL). Successful integration was positively selected for using G418 (GoldBio Cat no. G-418-5); animals retaining an extrachromosomal array were killed by heat-shock (2 hours, 34°C) induced *peel-1* toxicity. Genomic DNA was extracted from transgenic lines (QIAGEN Gentra Puregene Tissue Kit) and insertions were mapped by DpnII digestion of purified DNA, followed by ligation, inverse PCR, and Sanger sequencing.

### Cilia Imaging and Analysis

Imaging of ciliated sensory neurons was performed on adult males 24 hours post L4 stage. *C. elegans* were immobilized with 10 mM levamisole (Sigma) on 3% agarose pads. *clhm-1* expression pattern images were obtained with a Zeiss LSM880 confocal microscope. Images of splayed male tails for cilia analyses were acquired as Z stacks using a Zeiss LSM880 (63x oil objective) with Airyscan GaAsP-PMT area detector. Representative cilia images were acquired with an Andor Dragonfly microscope and Zyla sCMOS camera.

ImageJ (NIH) was used to plot normalized fluorescence intensity distribution. Starting beyond the distal tip, a linear ROI was drawn through the entirety of each individual cilium for RnB2-RnB5 neurons. The “multi plot” function was used to collect fluorescence intensity distribution profiles that were normalized individually to the maximum ROI value, then manually aligned using Microsoft Excel. Volocity (PerkinElmer) was used for quantitative volumetric analysis of ciliary colocalization and fluorescence signal intensity in three-dimensional reconstructions. Manders Correlation Coefficients (M1 and M2) were determined by drawing an ROI around each individual cilium and ciliary base. A high Manders coefficient indicates high correlation within a particular area; analysis of the relationship between M1 and M2 indicates pixel spread.

### EV Imaging and Analysis

Eight transgenic L4 hermaphrodites carrying the *him-5(e1490)* mutation were picked to 6 cm NGM plates and allowed to grow for 4 days, resulting in a mixed population of adult males and hermaphrodites. For analysis of EV release from virgin males, 20-30 L4 males were placed on a new plate on the third day of culturing and imaged the following day. Prior to imaging, adult males were picked to an unseeded plate, allowed to crawl for several seconds to clear bacteria, picked into 20 mM levamisole (100 mM diluted in Image-iT FX Signal Enhancer medium; ThermoFisher Item no. I36933) on 3% agarose pads, and covered with high-performance cover glass (Zeiss, item no.: 474030-9020-000).

Spectral imaging of EVs was performed with a Zeiss LSM880 confocal microscope using the standard spectral detector. ZEN black (Zeiss) was used for linear unmixing of single EV emission spectra. TIRF images of EVs were acquired with an Andor Dragonfly super-resolution microscope and Andor Zyla sCMOS detector. The TIRF angle of incidence was manually adjusted for each animal to achieve critical angle. All EV images were taken within 30 minutes of animal mounting.

Imaris software (Oxford Instruments) was used for quantitative EV analysis. EVs were identified using the “Spot” function, setting approximate object size to 0.350 µm in diameter and a quality threshold of 4 (GFP) and 10 (RFP), determined by analysis of negative controls (Fig. S2). Hot pixels and spots that intersected with the animal cuticle were manually removed. Intersecting vesicles were identified as GFP and RFP spots with a maximum distance of 0.3 µm.

### Statistical analysis and graphing

Dataset normality was determined using the Anderson-Darling normality test. Depending on normality, either the Student’s t-test or Mann-Whitney U test was used when comparing two data sets and one-way ANOVA or Kruskal-Wallis test with multiple comparisons when comparing three or more datasets. Statistical analyses and graphing were performed with GraphPad Prism 9; *p < 0.05, **p < 0.01, ***p < 0.001.

