## Supplemental Figures - Clupper et al for "Kinesin-II motors differentially impact biogenesis of distinct extracellular vesicle subpopulations shed from *C. elegans* sensory cilia"

### *Supplemental Figure 1*

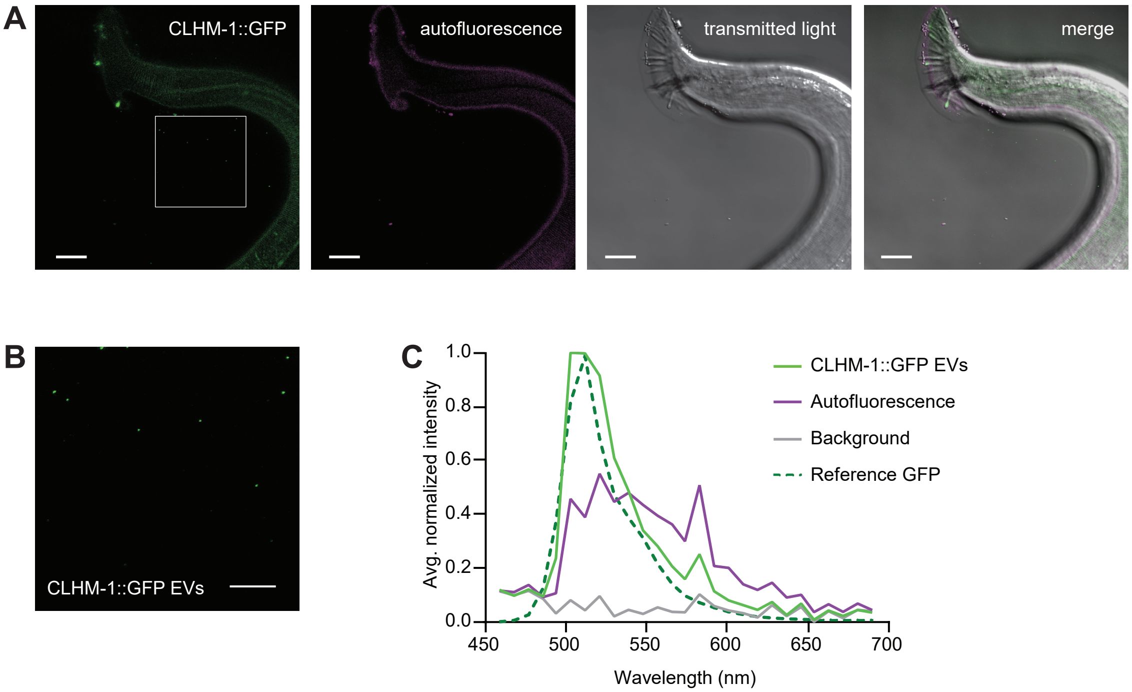

#### *Supplemental Figure 2*

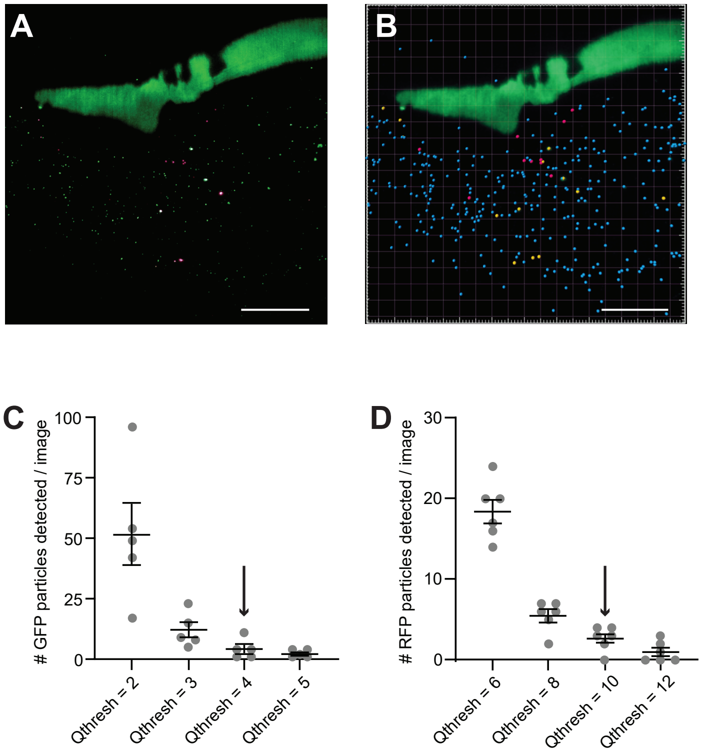

#### *Supplemental Figure 3*

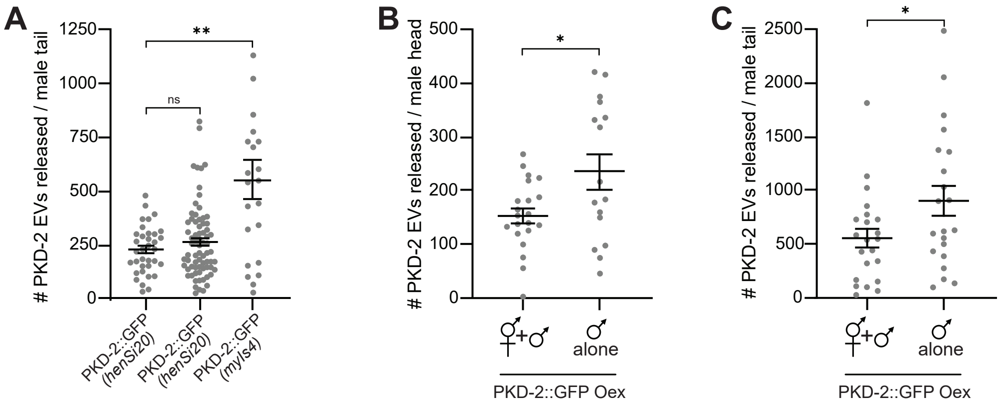

#### *Supplemental Figure 4*

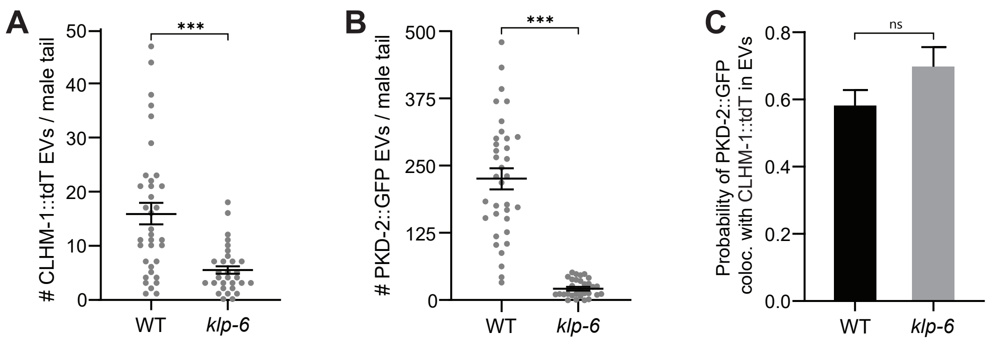

#### *Supplemental Figure 5*

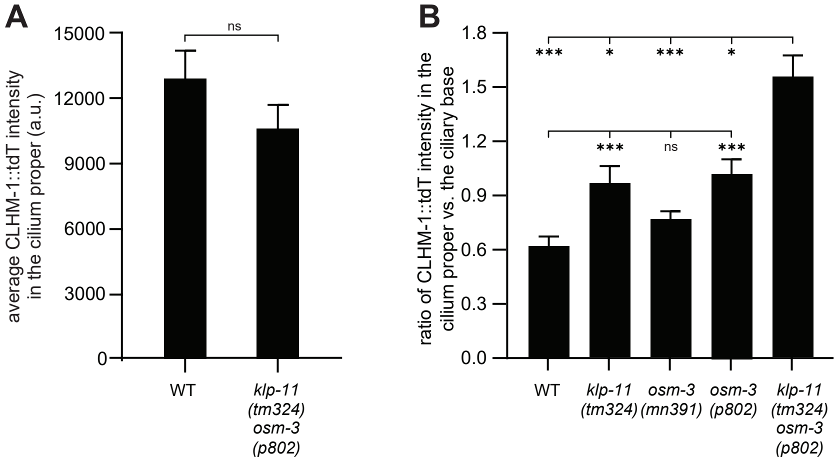

**Fig. S1 Fluorescence emission from CLHM-1::GFP EVs matches GFP reference spectrum**

A) Lambda imaging followed by linear unmixing allows for delineation between GFP fluorescence emission and autofluorescence in adult males expressing CLHM-1::GFP. Scale bars, 20 μm.

B) EVs containing CLHM-1::GFP (box, panel A); scale bar, 10 μm.

C) Fluorescence emission from linear unmixed channels. CLHM-1::GFP EV fluorescence emission matches the eGFP reference spectrum, not the autofluorescence spectrum or background noise.

**Fig. S2 Quantification of fluorescently-labeled environmental EVs using the Imaris spot detector function**

A) TIRF-acquired image of EVs released from the tail of a male expressing PKD-2::GFP and CLHM-1::tdTomato. EV imaging was performed across three platforms equipped with super-resolution capabilities: TIRF microscopy using the Andor Dragonfly (98 μm x 98 μm with 2048 x 2048 pixel resolution), Zeiss LSM880 confocal microscopy with Airyscan detection (73.95 x 73.95 μm with 1740 x 1740 pixel resolution), and structured illumination microscopy with the Zeiss Elyra PS1. All systems allowed for detection of fluorescent EVs, however, the Andor Dragonfly coupled with a Zyla sCMOS detector provided the largest field of view and greatest resolution. This platform allowed us to visualize the majority of EVs released per animal, resulting in robust and highly reproducible EV quantitation between different days and experimentalists.

B) Image from (A) showing fluorescently-tagged EVs identified with Imaris spot detection. Cyan spots label EVs containing PKD-2::GFP, magenta spots label EVs containing CLHM-1::tdTomato, and yellow spots label EVs containing both cargoes.

C) Average number of spots detected in the GFP channel for different quality thresholds in images taken of animals lacking a GFP transgene. At quality threshold 4 (arrow), detection of noise is sufficiently suppressed while detection of legitimate EV signal is minimally affected. Error bars show SEM; n = 6.

D) Average number of spots detected in the tdTomato channel for different quality thresholds in images taken of animals lacking a tdTomato transgene. At quality threshold 10 (arrow), detection of noise is sufficiently suppressed while detection of legitimate EV signal is minimally affected. Error bars show SEM; n = 5.

**Fig. S3 PKD-2 overexpression does not affect relative change in PKD-2 EV abundance between virgin and mated adult males**

A) Overexpression of PKD-2::GFP causes a significant increase in the detected number of EVs released from the male tail compared to males expressing single copy PKD-2::GFP transgenes; n ≥ 22.

B-C) Average number of PKD-2::GFP EVs released per virgin adult male head (B) and tail (C) is significantly higher compared to males raised with mating partners in the PKD-2::GFP overexpression strain; n ≥ 21.

Error bars show SEM; Kruskal-Wallis test used for (A), Student's t-test used for (B-C), * = $p<0.05$.

**Fig. S4 KLP-6 is required for release of both CLHM-1 and PKD-2 containing EVs**

A) *klp-6(sy511)* significantly reduced release of CLHM-1::tdTomato-containing EVs from male tail EVNs; n = 30.

B) Release of EVs containing PKD-2::GFP is significantly lower in *klp-6(sy511)* compared to wild type; n = 30.

C) Probability of PKD-2::GFP being present in a CLHM-1::tdTomato-containing EV is unchanged between wild type and *klp-6(sy511)*; n = 28.

Error bars show SEM; Mann-Whitney test, *** = $p<0.001$.

**Fig. S5 Loss of *klp-11* and *osm-3* reduces CLHM-1::tdTomato abundance in the ciliary base, but not the cilium proper**

A) CLHM-1::tdTomato fluorescence intensity in the RnB cilium proper does not change in *klp-11(tm324)*; *osm-3(p802)* double mutants compared to wild type n ≥ 16.

B) The ratio of CLHM-1::tdTomato fluorescence intensity in the cilium proper versus the ciliary base is greater in *klp-11(tm324)* and *osm-3(p802)* mutants compared to wild type. The *klp-11*;*osm-3* double mutant exhibited a further increase in the cilium proper to base fluorescence ratio, which is significantly higher than the wild type or any single mutant; $n \geq 16$.

Error bars show SEM; Kruskal-Wallis test, * = $p<0.05$, *** = $p<0.001$.
